# Pushing the limits of detection of weak binding using fragment based drug discovery: identification of new cyclophilin binders

**DOI:** 10.1101/136101

**Authors:** Charis Georgiou, Iain McNae, Martin Wear, Harris Ioannidis, Julien Michel, Malcolm Walkinshaw

**Author notes:** Corresponding Authors: Malcolm Walkinshaw, (for biophysical experiments), Julien Michel, (for computational chemistry).

## Abstract

Fragment Based Drug Discovery (FBDD) is an increasingly popular method to identify novel small-molecule drug candidates. One of the limitations of the approach is the difficulty of accurately characterizing weak binding events. This work reports a combination of X-ray diffraction, surface plasmon resonance (SPR) experiments and molecular dynamics (MD) simulations, for the characterisation of binders to different isoforms of the cyclophilin (Cyp) protein family. Although several Cyp inhibitors have been reported in the literature, it has proven challenging to achieve high binding selectivity for different isoforms of this protein family. The present studies have led to the identification of several structurally novel fragments that bind to diverse Cyp isoforms in distinct pockets with low millimolar dissociation constants. A detailed comparison of the merits and drawbacks of the experimental and computational techniques is presented, and emerging strategies for designing ligands with enhanced isoform specificity are described.

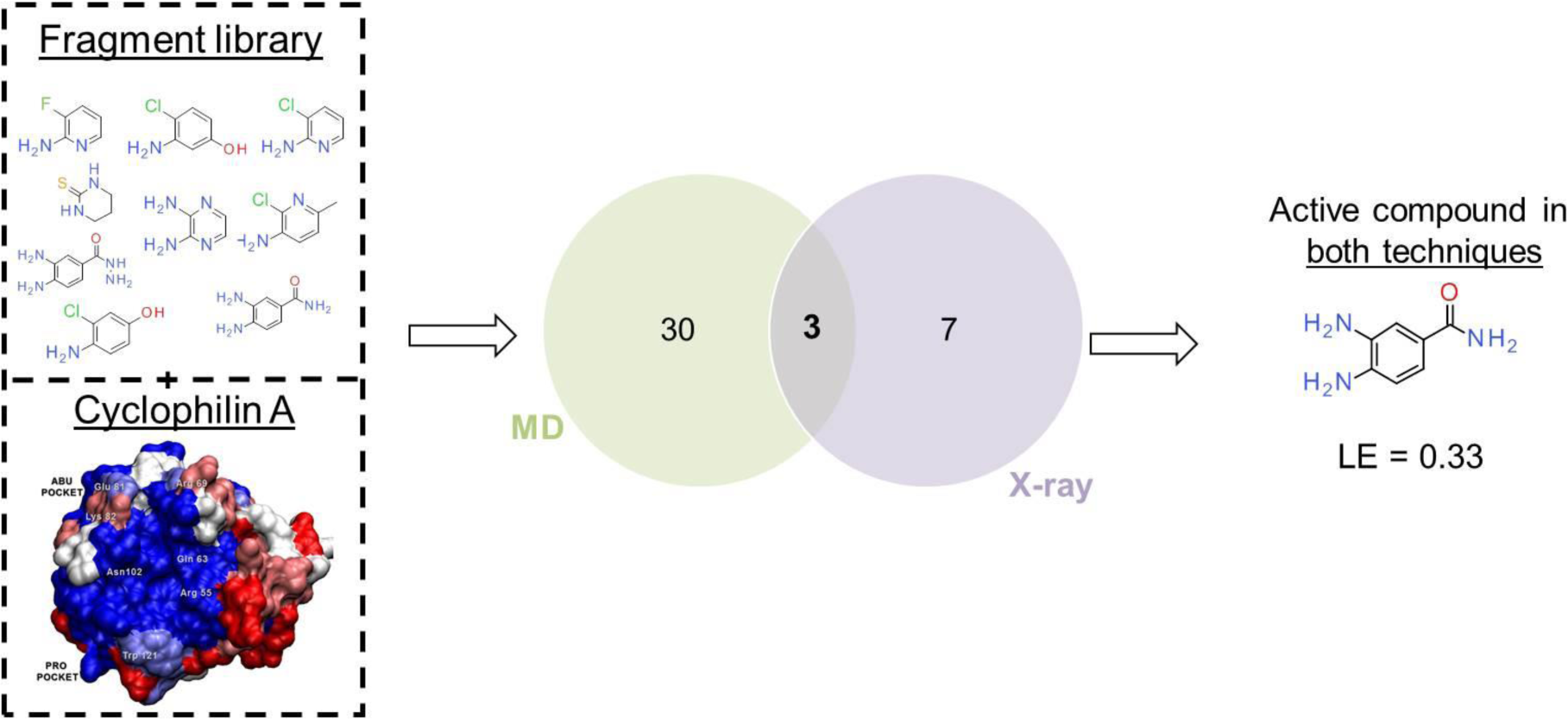

**Research Highlights:**

- FBDD is a popular method but weak binding is difficult to detect
- There is a need for pushing the limits of weak binding detection
- Combination of X-ray, SPR and MD methodologies increases successful characterization of weak binding events
- Several novel Cyclophilin fragment binders were identified

## Introduction

Fragment based drug discovery (FBDD) is frequently used to identify small organic molecules (fragments) as starting points for further structure-based drug design (SBDD) programs that aim to deliver drug-like molecules suitable for clinical studies. Fragments can be described using the “rule of 3” [1],[2,3]. According to this rule, a fragment is typically an organic molecule with molecular weight (Mw) ≤ 300 Da, number of H-bond acceptors ≤ 3, number of H-bond donors ≤ 3 and clogP (computed partition coefficient) ≤ 3 [1]. Fragments typically exhibit a dissociation constant in the micromolar to low millimolar range and the success of FBDD can be linked to steady improvements in robust biophysical characterisation of weak binding [2,4].

A number of problems are associated with the effective screening of fragment libraries. It is often difficult to solubilise fragments at concentrations required to saturate a protein target. Additionally the presence of aggregates, impurities and/or reactive intermediates can also lead to false positives or negatives. These issues are exacerbated for particularly small fragments (≤ 150 Da) that are likely to exhibit at best mM dissociation constants, and there is a need for pushing the limits of detection of weak binding to broaden the scope of FBDD.

The past decade has seen rapid developments in the application of molecular simulations to structure-based drug design [5]. Molecular simulations and free energy calculations are now being used to complement experimental approaches for a wide range of protein – ligand complexes, including estimation of binding energies of drug-like molecules [6,7], and fragments [8]. This report focuses on the combination of biophysical measurements and molecular simulation methods to characterize weak Cyclophilins (Cyps) binders present within a library of small fragments.

Cyclophilins (Cyps) are a family of peptidyl-prolyl isomerases (PPIases) that catalyze the isomerization of proline residues, promoting and facilitating protein folding [9]. The human cyclophilin family counts seventeen members with the prototype and most abundant being cyclophilin A (CypA) [10]. Cyclophilin orthologues can also be found in most plants, parasites and animals [11,12]. The amino acid sequences of all human Cyps are well established, as well as their secondary and tertiary structures [10]. All Cyps share a highly conserved PPIase domain. Cyps are also members of the immunophilins class of proteins [13–15] and many Cyps are inhibited by the natural immunosuppressant cyclosporin A (CsA) [10,15]. CsA is used in organ transplantation to prevent immune response and organ rejection. Cyps are also involved in (mis)regulation of several biological signaling pathways including damage-induced cell death [16], RNA splicing [17] and different types of cancer. Cyps are also known to be involved in the life cycle of different viruses such as Human Immunodeficiency Virus 1 (HIV-1) and Hepatitis C Virus (HCV) [18,19].

Because of their diverse biological roles Cyps are recognized as potential biological targets for the treatment of HCV [19–21], HIV [22–24], cancer [25–27] and neurodegenerative diseases such as Parkinson’s and Alzheimer’s [28–30]. Originally, efforts for the identification of cyclophilin inhibitors were focused on cyclic peptides, analogues of CsA [20,21,31,32]. In recent years a number of small-molecule Cyp inhibitors have been reported in the literature as [33–41]. A major unsolved challenge is to achieve robust binding affinity and specificity to distinct Cyp isoforms. Difficulties in achieving strong binding affinities arise from the shallow, solvent-exposed nature of the active site of Cyps. Further, the high degree of structural similarity between isoforms makes it challenging to achieve high binding specificity. Nevertheless this is widely thought to be necessary to produce chemical probes able to elucidate the biological roles of different Cyp isoforms, and to pave the way for next-generation Cyp drugs with reduced side-effects in comparison with CsA analogues.

The present work used Surface Plasmon Resonance, X-ray diffraction and molecular dynamics simulations to screen a focused library of small fragments against the most common Cyp isoforms Cyp A, B and D. This combination of multiple methodologies lead to the characterization of novel Cyp fragments that are suitable starting points for further optimization into more potent and isoform specific lead molecules.

## Results

### A focussed fragment library to interrogate ligandability of the Abu pocket in Cyclophilins

The starting point for this focused library design was 2,3-diaminopyridine, for which SPR and X-ray data had been generated previously, demonstrating the stoichiometric binding of this fragment to the Abu pocket of CypA (Figure 1) [42]. Comparative analysis of primary protein sequences has suggested that the Abu pocket in Cyps may be utilised to engineer isoform-selective ligands owing to small variations in amino-acids that line up the edge of the pocket between the common isoforms Cyp A-D. This contrasts with the nearby Pro pocket that is more conserved across the Cyp family. Thus small fragment analogues of 2,3-diaminopyridine were selected to study the chemical diversity that would be tolerated by the Abu pocket. A total of one hundred small fragments, structurally distinct from those previously tested, were chosen based on chemical similarity and commercial availability. Analogues include substituted aromatic rings such as pyridines, pyrazines, pyrimidines, as well as non-aromatic rings. A full list is provided in the supplementary Table 1. The relatively low average molecular weight (ca. 150 g mol^−1^) of the library members is dictated by the small size of the Abu pocket, hence any binders are expected to exhibit at best high micromolar to low millimolar binding constants.

**Fig.1.**
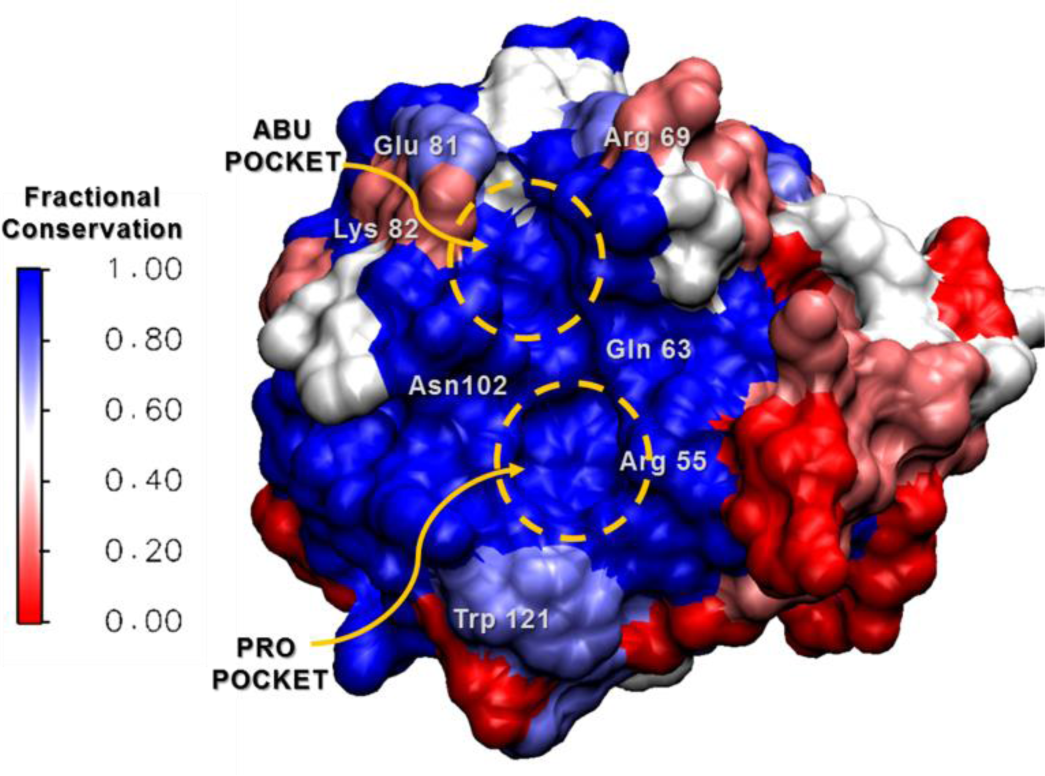
3D surface structure of CypA colour coded according to degree of residue conservation across isoforms A, B, C, D, F (blue – strictly conserved, red – different in every isoform). Abu and Pro pockets and key active side residues are also highlighted.

### SPR suggests several of the fragments may bind weakly to diverse Cyp isoforms

All the fragments were initially screened by surface plasmon resonance at 1 mM, using conditions described previously [42], on a high-density (to account for the mass ratio of the fragments to His-CypA, -B and -D) surface of 3200, 3000 and 3000 RU covalently stabilized His-CypA, -B and –D, respectively. Apparent Cyp specific hits were further analysed with a 2-fold concentration series from 0.015 mM to 1 mM.

This exercise provided evidence for specific binding in the range of 0.5 – 20 mM against at least one isoform for approximately 15 fragments. However it was not possible to derive reliable *K_d_* estimates due to limitations in compounds solubility, and difficulties in rigorously removing potential refractive index artefacts at the analyte concentrations tested. Thus the outcome of the SPR screen was deemed overall encouraging, but further characterisation of potential binders was pursued via other means.

### Free energy calculations estimate fragment binding energies in line with experimental data, and suggest binding pocket preferences

The standard (absolute) binding free energies of 86 of the 100 experimentally tested fragments were estimated by alchemical free energy calculations.[43] The 14 excluded compounds were either carrying out a net charge, or were too structurally dissimilar to other compounds in the library to allow straightforward computation of relative free energies of binding (see methods). The overall objective of the project was to discover new Abu pocket binders, but fragments could in principle bind to either the Abu or Pro pockets. Therefore two sets of calculations were executed with fragments docked in the Abu or Pro pockets respectively. Figure 2 depicts a histogram that summarizes the results of these calculations.

**Fig.2.**
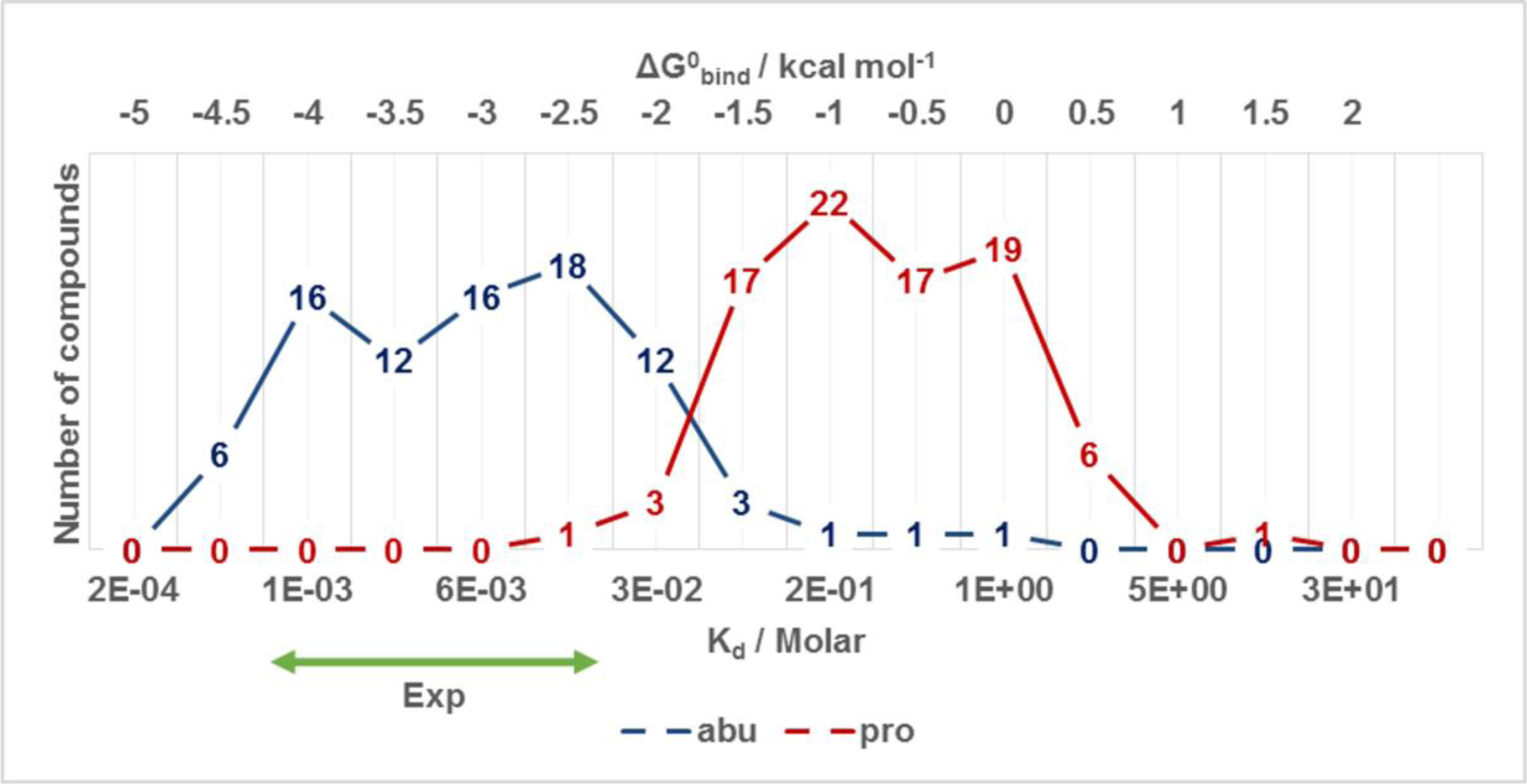
Histogram of calculated binding free energy of fragments to Abu and Pro pockets of CypA. Compounds are binned (bin width 0.5 kcal mol^−1^) based on their calculated *K_d_* (bottom *x*-axis) and calculated *ΔG*°*_bind_* values (top *x*-axis). The coloured numbers indicate the total number of compounds in each bin. The calculated binding energies for the Abu and Pro pockets are depicted in blue and red respectively. The SPR range of *K_d_* estimates for fragments bound to CypA is depicted by the green arrow.

The data shows that there is a clear preference for the fragments to bind to the Abu pocket of Cyp A, with a mean and standard deviation of binding energies of -3.0±0.7 kcal.mol^−1^ and -0.7±0.5 kcal.mol^−1^ for the Abu and Pro pockets respectively. This was deemed encouraging and reflective of the strategy used to assemble a focused library targeting the Abu pocket. Conversion of the calculated absolute binding free energies into dissociation constants indicate that the range of predicted *K_d_*s to the Abu pocket of CypA is between 0.5 – 30 mM, whereas for the Pro pocket the range is 30 mM – 2 mol L^−1^. The computed binding energies preferences for the Abu pocket were in line with the 0.5 – 20 mM binding constant range that was estimated from the SPR screen. Altogether the molecular dynamics data was supportive of preferential weak mM binding to the Abu pocket for several fragments in the focussed library.

### X-ray crystallography provides evidence of binding for several fragments

A total of 40 compounds were selected for further X-ray crystallographic studies on CypA, CypB and CypD. The selection of this subset was guided by a combination of factors including: evidence of binding to a Cyp surface in the preceding SPR screen; computed interactions with the Abu/Pro pockets; desire for producing co-crystal structures of fragments bounds to multiple isoforms; evidence of sufficient solubility; available beam-time.

Unfortunately no crystal structures of fragments in complex with CypB or CypD were obtained. Although well diffracting crystals were obtained for CypB, the crystal packing arrangement blocked access to the active site and soaking experiments were thus not successful. CypD yielded crystals that were too fragile for soaking and data collection.

Over thirty compounds that were soaked into CypA crystals diffracted well. Refinement of the structural data led to the detection of 10 fragments in complex with CypA. The chemical structures of all such fragments are given in Table 1 and data collection, refinement and Ramachandran plot statistics for each of the CypA complexes are reported in the supplementary Table 2. Fragments that were not found in the active site of CypA are listed in the supplementary Table 3. The majority of the observed compounds bind in the Abu pocket, in line with the expectations from the library design strategy and MD calculations, however some fragments were found to bind to the neighbouring Pro pocket. The pocket preference appears to be dictated by the nature of H-bond donors and acceptors, and also the hydrophobicity of each ligand.

**Table 1.**
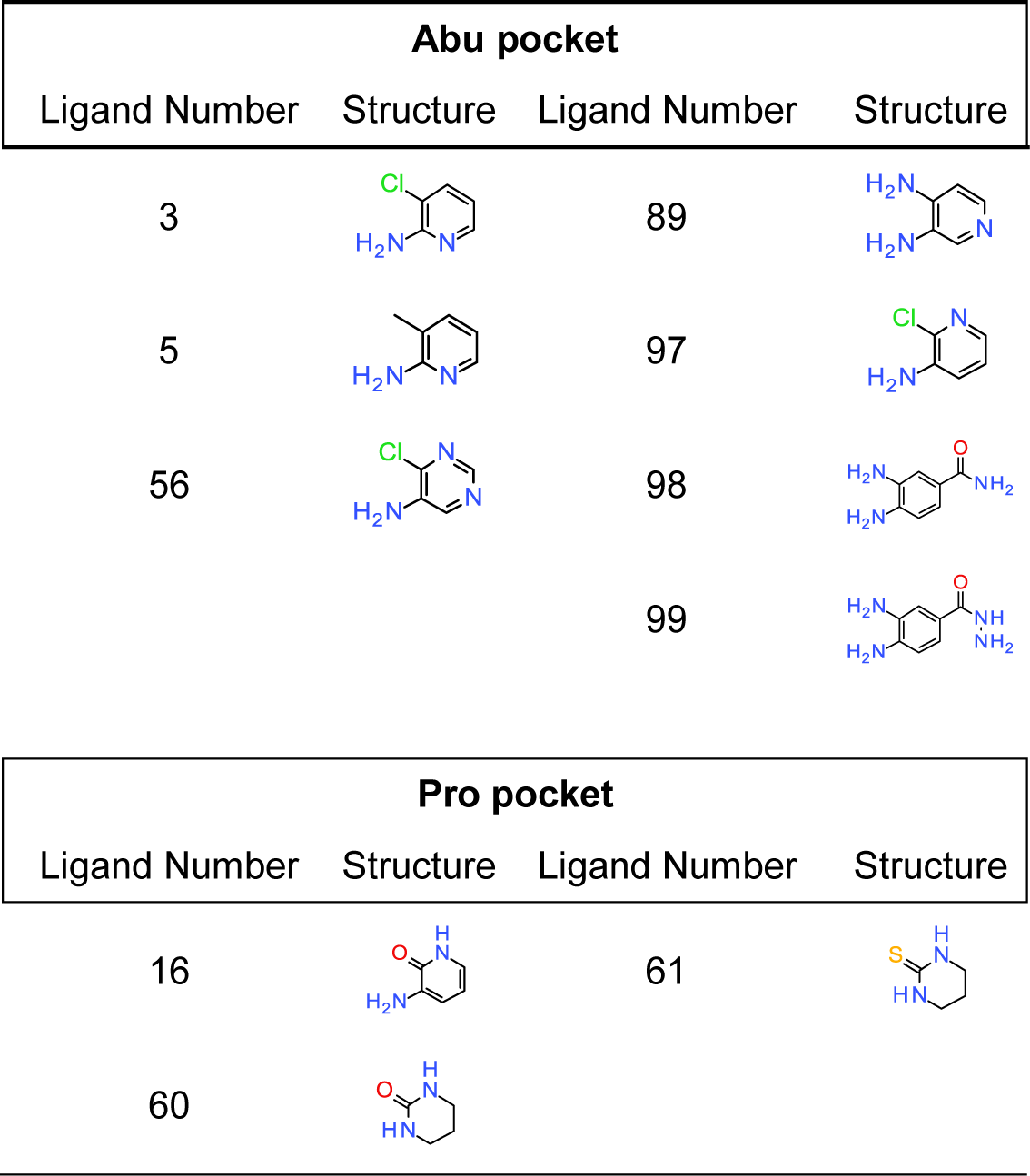
Fragments that bind to CypA based on X-ray crystallography data.

| Abu pocket |  |  |  |
| --- | --- | --- | --- |
| Ligand Number | Structure | Ligand Number | Structure |
| 3 |  | 89 |  |
| 5 |  | 97 |  |
| 56 |  | 98 |  |
|  |  | 99 |  |

| Pro pocket |  |  |  |
| --- | --- | --- | --- |
| Ligand Number | Structure | Ligand Number | Structure |
| 16 |  | 61 |  |
| 60 |  |  |  |

Ligands with hydrogen bond donors and acceptors such as **3**, **5**, **56**, **89**, **97**, **98** and **99** bind to the more hydrophilic Abu pocket of CypA. This is in agreement with the MD simulations that predicted a preference for the Abu over the Pro pocket for all 7 fragments (mean preference -2.6±1.0 kcal.mol^−1^). These compounds adopt broadly similar binding poses and interactions with the Abu pocket. Compounds **89**, **98** and **99** have a very similar binding pose, and the electron density is well defined (Supplementary Fig. 2). Figure 3 depicts fragment **98** and shows how the 4-amino group forms hydrogen bonds with two water molecules, W1 (violet-purple) and W2 (sky-blue). Both W1 and W2 have B-factors values (ca. 15 Å^2^) lower than for most other water molecules present in the crystal structure. This was taken as indication that these two hydration sites are well ordered. By contrast the 3-amino group is H-bonded to two additional mobile water molecules W3 (pink) and W4 (green). These water molecules have higher B-factors values (ca. 30 Å^2^) and are thus expected to be less ordered. Overall this somewhat unusual cluster of water mediated protein-fragment interactions play a key role for fragment binding to the Abu pocket. W1 forms H-bond interactions with three protein residues; Ala101, Gln111 and Gly109; W2 with Gly74, Ser110 and Gln111; W3 with Glu81, Gly75 and W4 with Thr73.

**Fig.3.**
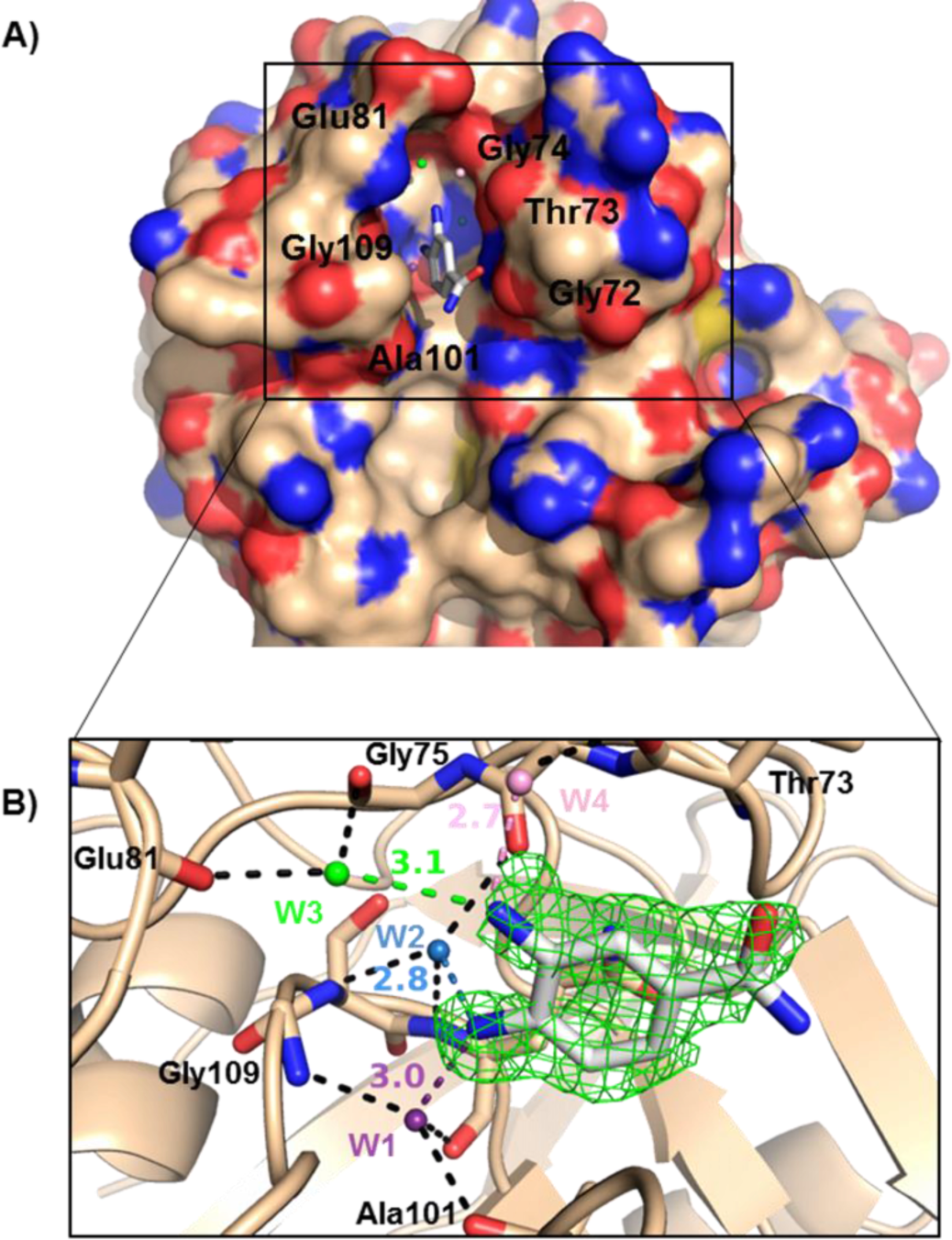
Binding of **98** in the Abu pocket of CypA. **A)** 3D surface structure of **98** in the Abu pocket of CypA. **B)** The inset shows a *Fo* – *Fc* omit map of **98** in the Abu pocket CypA. A 3*σ* contour is shown in green. H-bonds between ligand and water molecules are highlighted and colour coded as; W1 in violet-purple, W2 in sky blue W3 in green, W4 in light pink and W5 in orange (distances in Angstrom). H-bond interactions between water molecules and CypA residues are also highlighted in black.

Additionally a diverse range of small functional groups (halides, amino, methyl) is tolerated at a third position on the aromatic ring. Fragments **3, 5, 56** and **97** also bind in the Abu pocket of CypA and occupy the same position as **89**, **98** and **99**, though their electron density is less well defined. An unambiguous assignment of their binding poses is not possible and, in light of the molecular dynamics simulation results (see below), the electron density map was interpreted as the result of these ligands adopting multiple poses.

Fragment **56** seems to be able to adopt a slightly different orientation in the Abu pocket, with the chloro group of the ring buried in the pocket, whereas the amino group is more water exposed. This alternative pose is stabilised by two other water molecules that interact with the amine and the pyrimidine nitrogen atoms of the fragments and residues Gly63 and Thr73. The structure of all compounds bound to the Abu pocket, as well as their electron density and distances between fragments and protein/water molecules can be seen in Supplementary Fig. 2.

Some fragments were found to also bind out of a pocket in the vicinity of the 80’s loop. Figure 4 shows fragment **3** as an example. This second binding site lies between two symmetry related Cyp molecules in crystal CypA with halogen – carbonyl contacts to Asn106 and Glu84 and an aromatic interaction with Trp121 of another CypA molecule. This site was deemed to be a crystallographic artefact and ignored for further studies.

**Fig.4.**
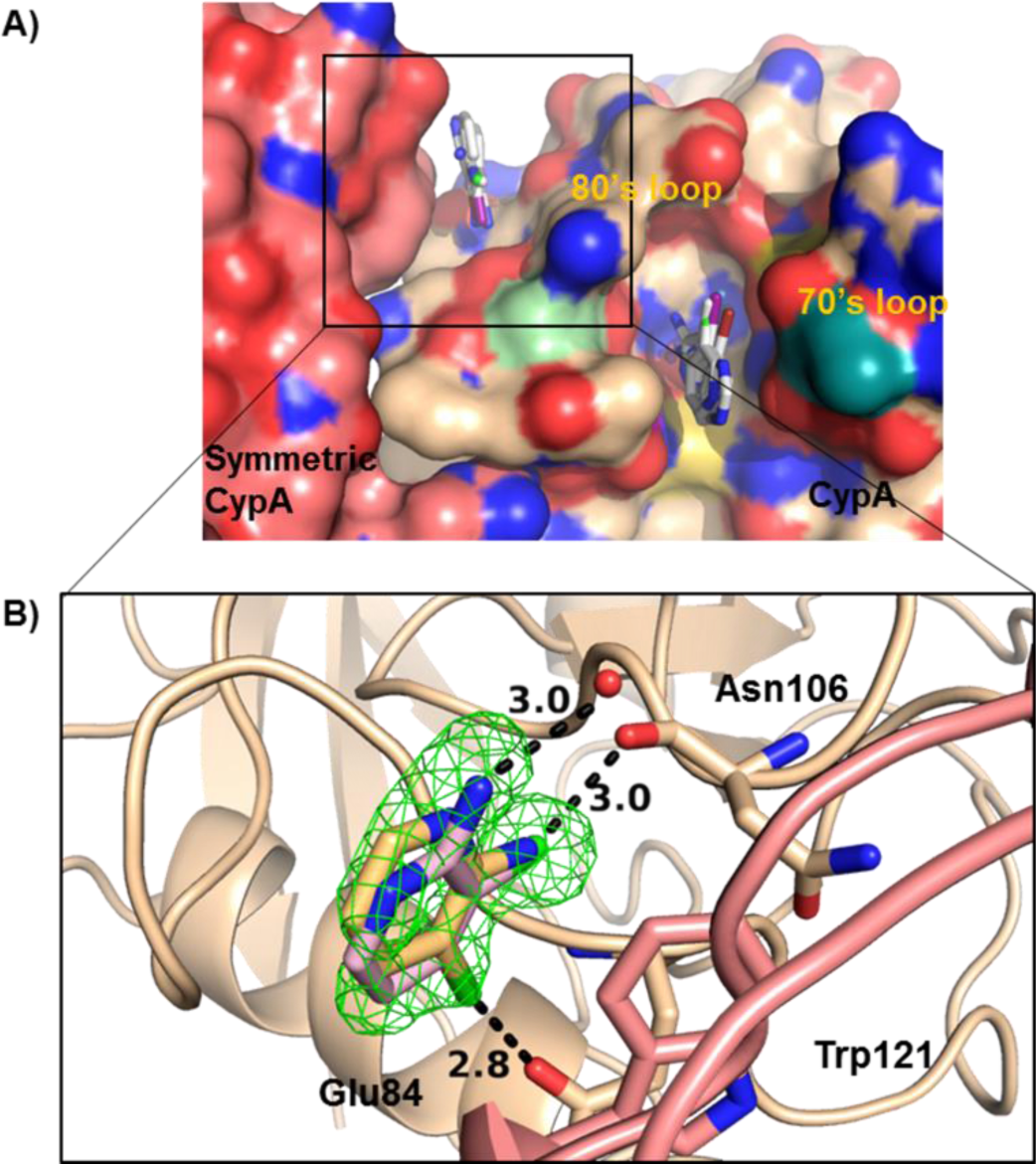
Binding of fragments out of the 80’s loop of CypA. **A)** Overlay of **3**, **5**, **56** and **97** bound to a 3D surface representation of CypA. A symmetric CypA structure can be seen in pink. **B)** The inset shows interactions between two alternative conformations of **3** (light orange and light pink) with Asn106 and Glu84 residues of CypA and a close van der Waals interactions with Trp121 in the symmetric CypA unit. A *Fo* – *Fc* omit map of **3** contoured at 3*σ* is shown in green.

Three of the ten fragments, **16**, **60** and **61** were found to bind the Pro pocket of CypA instead of the Abu pocket. This contrasts with the MD simulations that predicted a preference for binding to the Abu pocket (mean preference -1.7±0.7 kcal.mol^−1^). Figure 5 shows that crystallography indicates the three fragments establish H-bond interactions in the Pro pocket with the conserved CypA residues, Arg55, Gln63 and Asn102. In contrast to the Abu pocket binders that all contain one aromatic ring, fragments **16**, **60**, **61** are made of non-planar six membered rings. Thus a combination of lack of planarity and increased hydrophobicity seems to bias fragments towards binding to the conserved Pro pocket.

**Fig.5.**
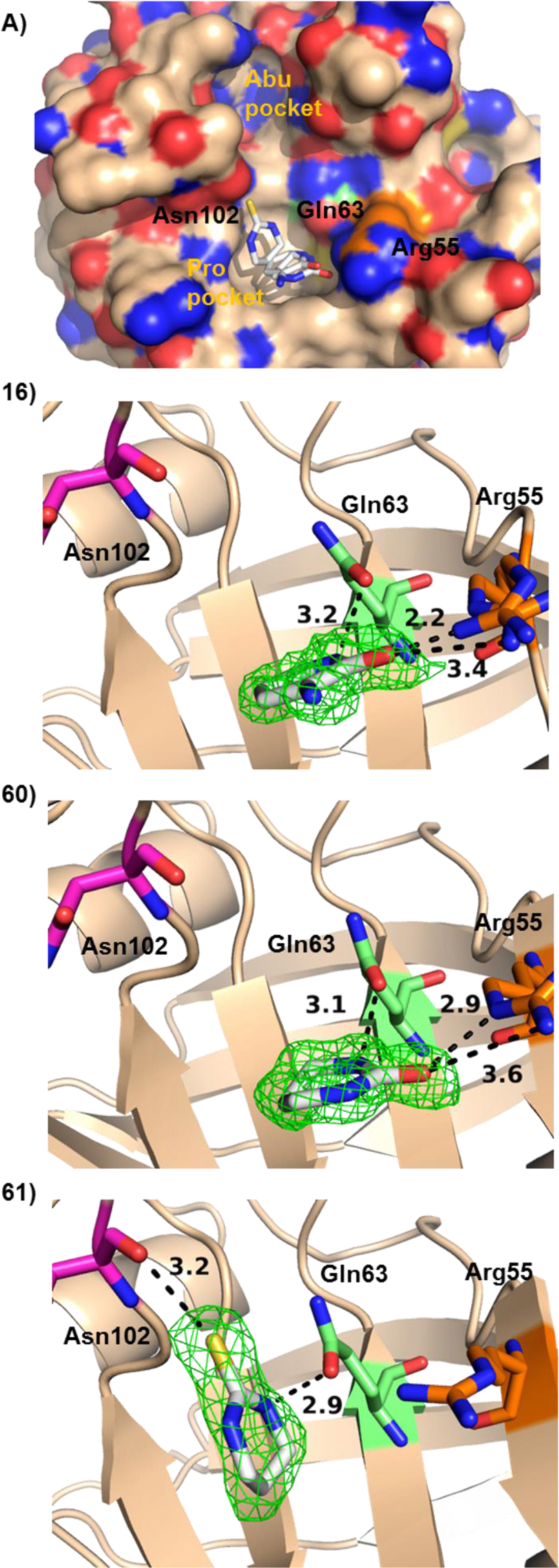
Bound poses of **16**, **60** and **61** in the Pro pocket of CypA. **A)** Overlay of **16**, **60** and **16** bound to Cyp A (3D surface representation). The three other panels depict interactions of **16**, **60** and **61** with Arg55, Gln63 and Asn102 residues of CypA in the Pro pocket. *Fo* – *Fc* omit maps contoured at 3*σ* are shown in green. Distances between ligands and CypA residues are highlighted in black and given in Angstrom.

### Molecular dynamics simulations suggest that some fragments adopt multiple binding modes and help guide the interpretation of X-ray diffraction measurements

The structures of fragments derived by X-ray crystallographic analyses revealed a conserved binding mode that appears to be adopted by all fragments binding to the Abu pocket (Fig. 3). However for some fragments (e.g. **3**, **5, 56** and **97**) refinement of the structures was ambiguous and suggested that these fragments may adopt two different binding modes in the Abu pocket. This was investigated further by carrying out a detailed analysis of MD simulations of **3** bound to Cyp A. Since **3** shows a high degree of structural similarity to the other fragments found to the Abu pocket, it is anticipated that the observed interactions are also relevant for the other fragments. Figure 6 shows RMSD distributions of **3** from one X-ray crystallographic pose calculated from the MD simulations. Supplementary movie M1 depicts one representative MD simulated trajectory used for the RMSD analysis. A total of three different clusters are apparent in the distribution. Monitoring of hydrogen bonding distances between ligand/protein donor/acceptor groups also indicates three clusters whose structure and population match the results produced by RMSD analyses (Supplementary Fig. 3). The first cluster, denoted “MD pose A”, corresponds to a binding mode very similar to one X-ray pose (“X-ray pose A”) with a RMSD of 1 ± 1 Å. The second cluster, “MD pose B”, corresponds to a binding mode very similar to another X-ray pose (‘’X-ray pose B”) with a RMSD of 0.8± 0.3 Å. The last cluster (“MD pose C”) deviates more significantly from the X-ray data, with RMSD values of 3.8 ± 0.5 Å and 2.5 ± 0.5 Å to X-ray pose A and X-ray pose B respectively. MD pose A accounts for approximately 30% of the conformations, MD pose pose B accounts for 55% of the conformations, and MD pose C accounts for the remaining 15%.

**Fig.6.**
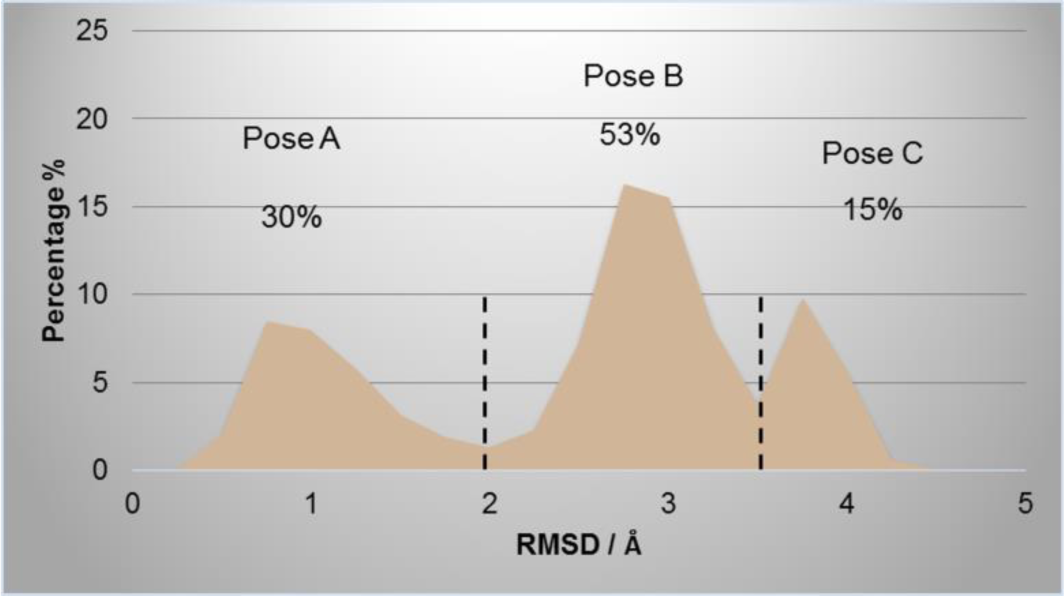
Histogram of RMSD of **3** from the X-ray crystallographic pose A computed from MD simulation trajectories.

Detailed visualisation of the MD trajectories indicates the interactions that account for the stability of each pose. Figure 7 shows four representative trajectory snapshots from MD pose A, indicating that compound **3** remains flexible, with its amino group interconverting rapidly between alternative hydrogen-bonding arrangements involving Thr107 (pale-green), Ala101 (light yellow), W1 (violet-purple) and W2 (sky blue). Figure 8a shows a representative snapshot from MD pose B. Compound **3** is rotated by 90° in the Abu pocket with respect to MD pose A, and is H-bonded to Thr73 (cyan) and W2 (orange). This is the main interaction pattern observed in MD pose B. Figure 8b depicts a representative trajectory snapshot of compound **3** in MD pose C. This pose is further rotated by 90° degrees with respect to pose B to position the amino group of **3** in the direction of the Pro pocket. This pose features mainly hydrogen bonding interactions with the backbone carbonyl oxygen of Gly72 (red). Detailed statistics about hydrogen-bonding interactions in each poses are given in supplementary figure 3 and 4.

**Fig.7.**
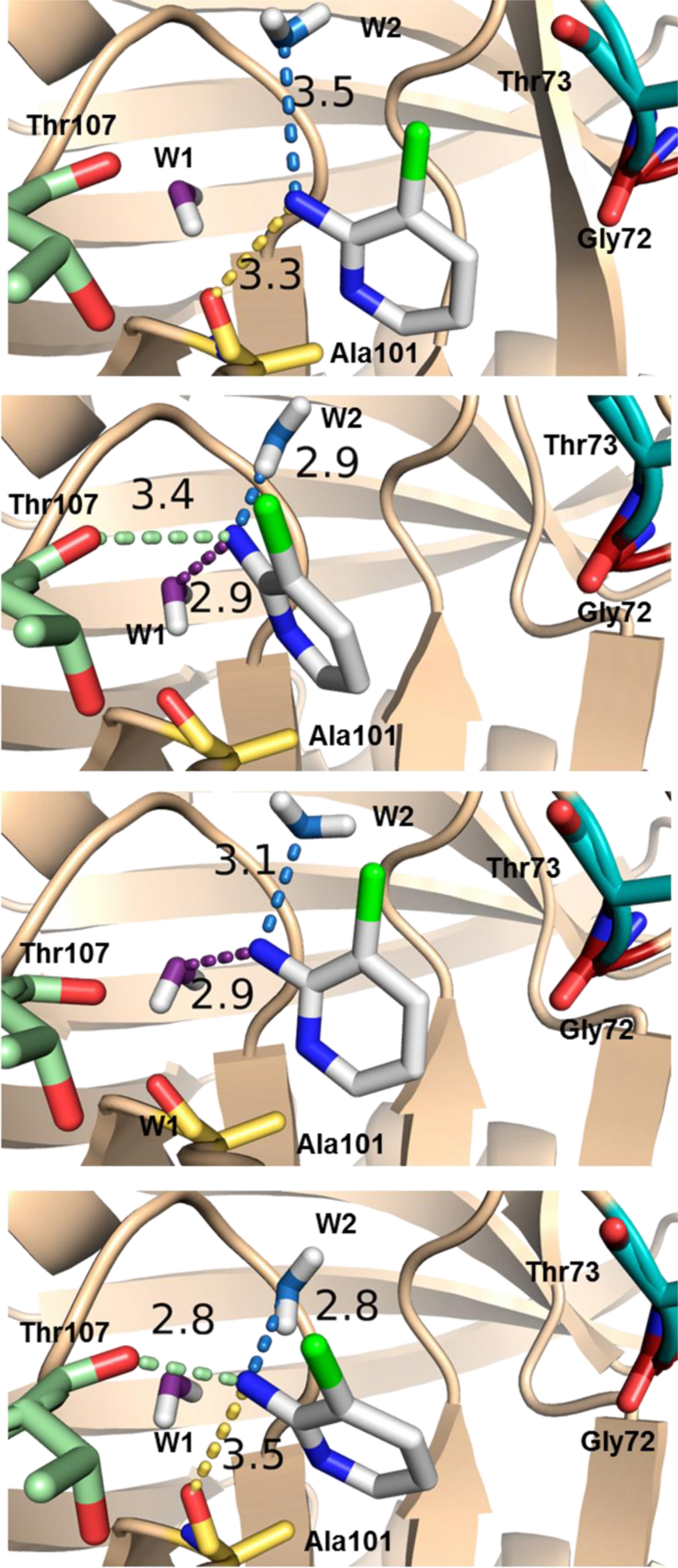
Representative conformations of ligand **3** in MD pose A sampled by the MD simulations. Protein residues or water molecules that are involved in H-bonding with **3** are colour coded separately. The distances between hydrogen bond donors and acceptors are given in Angstrom.

**Fig.8.**
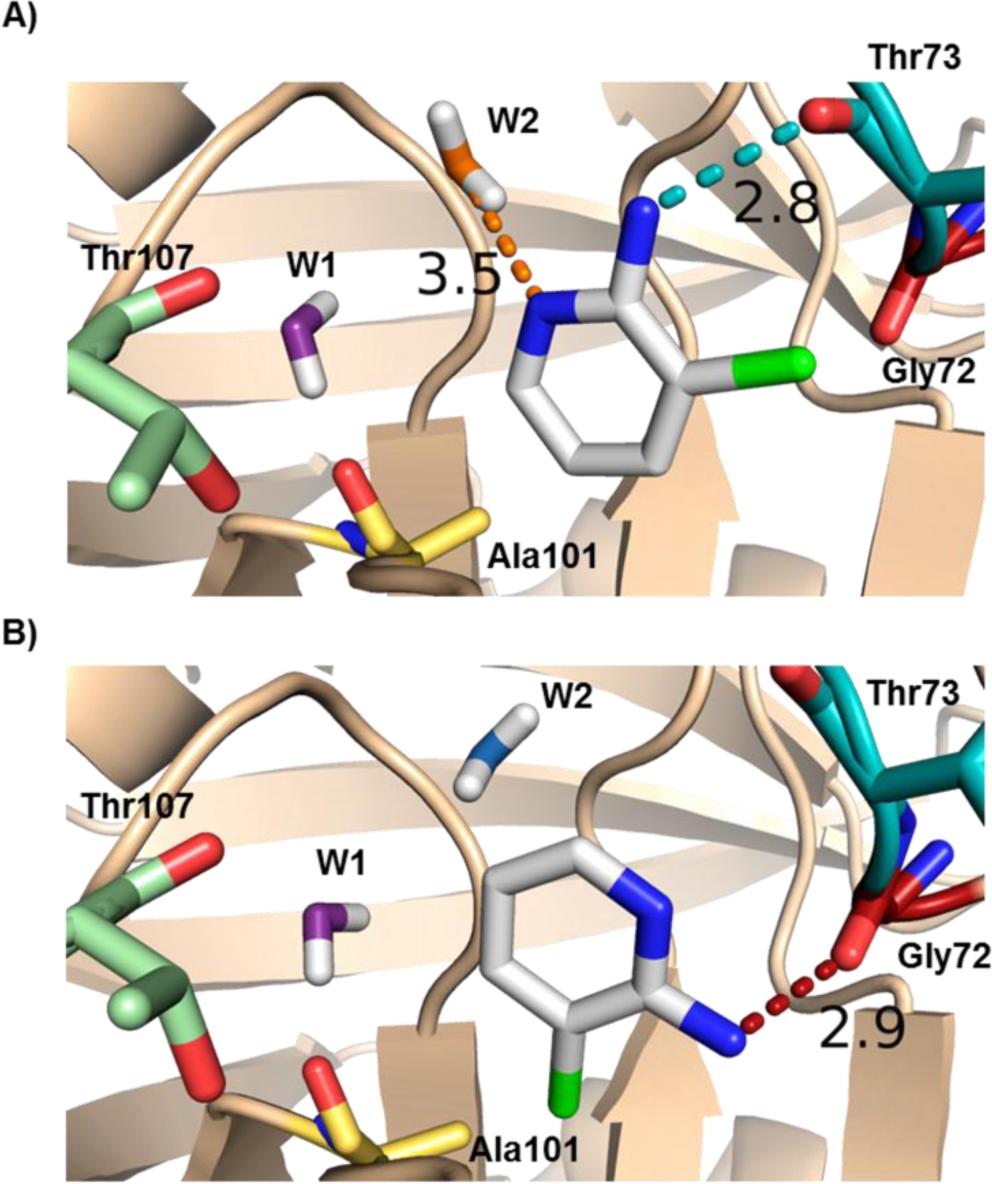
Representative conformation of ligand **3** in MD pose B (**A**) and MD pose C (**B**). Protein residues or water molecules that are involved in H-bonding with **3** are colour coded separately. The distances between hydrogen bond donors and acceptors are given in Angstrom.

In all MD trajectories water molecules W1 and W2 are very tightly bound in the Abu pocket and stay in the same orientation for the duration of the simulations. This is in agreement with the X-ray data which showed W1 and W2 present in the Abu pocket in all structures with low B-factors. In contrast to W1 and W2, water molecules W3 and W4 are more mobile, and easily displaced by other water molecules during the MD simulations. This is again in agreement with the X-ray results which showed W3 and W4 to have higher B-factor values compared to the W1 and W2 crystal water sites.

Overall this detailed analysis increased confidence in the interpretation of electron density maps derived from X-ray crystallography, and ligands **3** and **97** were ultimately refined in two different poses with 50% occupancy each (Supplementary Figure 2).

## Discussion and Conclusion

### How confidently can weak mM fragment binders be determined?

The combined data from SPR, X-ray crystallography measurements and MD simulations provides solid evidence that several of the fragments from the present in-house bespoke library interact with Cyp surfaces with dissociation constants in the low millimolar range. Figure 9 depicts a Venn diagram demonstrating overlap between ‘hits’ detected between X-ray analyses and MD simulations in the case of Cyp A. The SPR data was deemed too ambiguous to enable meaningful evaluation of hit rates and is thus not included in Figure 9. Given the criteria used here for declaring a compound a ‘hit’, X-ray and MD detected 10 and 33 binders respectively. This corresponds to a hit rate of 25-33% for X-ray (10 out of 40, or 10 out 30 if poorly diffracting crystals are excluded from the dataset), and 38% for MD (33 out of 86). For X-ray the definition of a hit is relatively clear (structure of a bound fragment refined against observed electron density), but many samples could not be analysed to detect ligands due to low quality diffraction data. Additionally, the methodology as used here doesn’t yield *K_d_* values and other techniques must be used to estimate binding constants for the fragments. By contrast for MD the criteria for declaring a hit is less clear cut since binding constant estimates are obtained for all compounds tested. Given the observed distribution of computed binding energies in Figure 2, a cutoff in standard binding free energies of -3.5 kcal.mol^−1^ was deemed reasonable. This corresponds to dissociation constants in line with the SPR estimates, and ligand efficiencies in the range of 0.3-0.5. Of course the number of hits can be decreased merely by setting more stringent requirements for the MD binding energy estimates.

**Fig.9.**
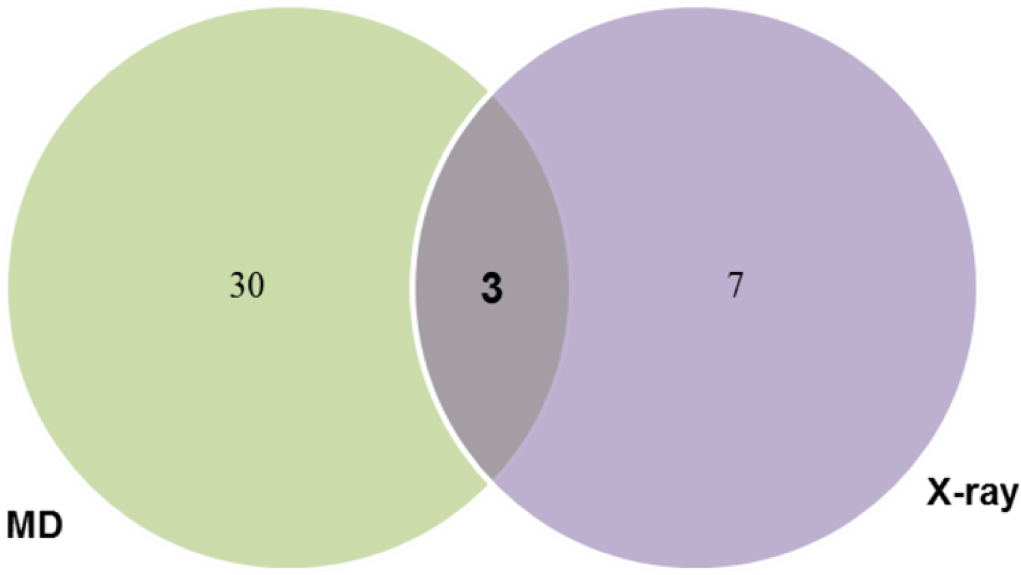
Venn diagram showing the number of hit compounds produced by X-ray crystallography and molecular dynamics techniques.

Overall 3 compounds were found active in both techniques. Out of these 3 binders, compound **98** was deemed of particular interest (Table 2). MD simulations suggest a CypA binding energy of -3.7 ± 0.4 kcal mol^−1^. The corresponding ligand efficiency is reasonable given the challenges posed by the Abu pocket in Cyclophilins. Compound **98** was also observed by X-ray crystallography in the Abu pocket of CypA, establishing H-bond interactions with water molecules and protein residues according to the canonical pattern observed for other Abu pocket fragments.

**Table 2.**
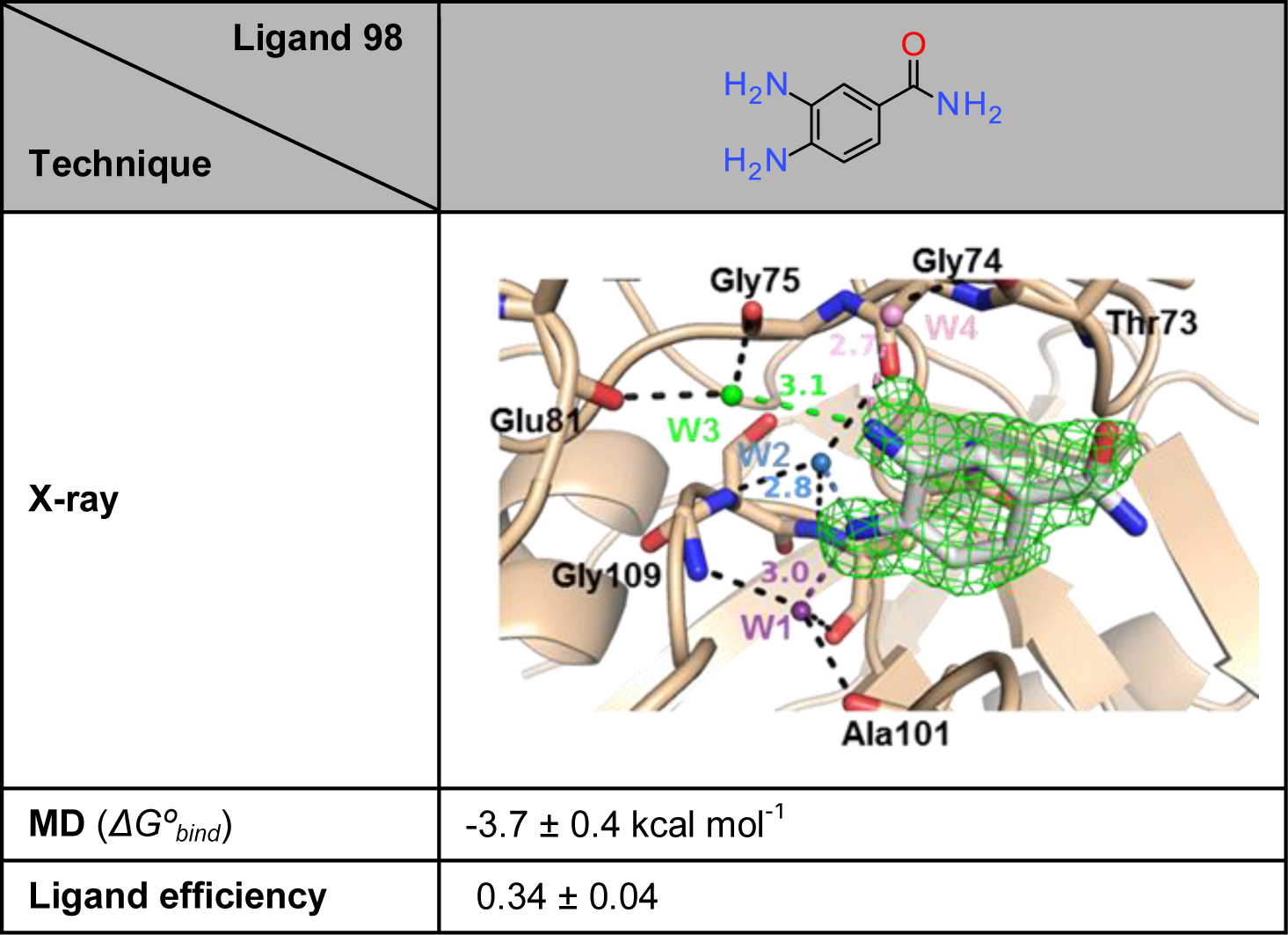
Summary of structural data for ligand **98**.

| Ligand 98<br>Technique |  |
| --- | --- |
| X-ray |  |
| MD ($\Delta G_{bind}^o$) | $-3.7 \pm 0.4 \text{ kcal mol}^{-1}$ |
| Ligand efficiency | $0.34 \pm 0.04$ |

### How can molecular dynamics complement biophysical methods in fragment-based drug discovery?

The binding free energies computed via molecular dynamics simulations were able to reproduce the preference for the binding of the fragments to the Abu pocket instead of the Pro pocket of Cyp A. Additionally the MD binding affinity estimates were in line with the range inferred from SPR analysis. These observations suggest that MD simulations can be used to guide the design of focussed fragment libraries enriched in structures more likely to bind a target pocket with a reasonable binding affinity. This contrasts with the more common strategy that involves assembling a generic library of structurally diverse fragments as a primary screening library.

Nevertheless a significant number of MD ‘hits’ were not observed via X-ray diffraction experiments. Arguably, reliably rank-ordering compounds by potency is particularly challenging for this dataset since by design the focussed library contains compounds that are structurally similar, and thus can all be reasonably modelled in the Abu pocket. Most of the library compounds show a computed standard binding energy that falls within the interval of -3 ± 1 kcal mol^−1^ (Figure 2). The relatively small dynamic range of calculated binding energies poses challenges given the precision of the free energy estimates (ca. 0.5 kcal mol^−1^), and the typical accuracy of classical biomolecular force-fields (typical systematic errors around 1 kcal mol^−1^ in favourable cases). It is likely that greater performance may be observed for the scoring of structurally diverse fragment libraries. This hypothesis could be tested by blinded-predictions, in a fashion similar to competitions for blinded estimations of standard binding energies of host/guest and protein/drug-like complexes have been organised recently. [44,45] It also cannot be ruled out that the conditions used to solve protein structures in X-ray crystallography experiments disfavour the binding of some of the fragments predicted by MD simulations to bind in solution.

Analysis and visualisation of MD trajectories for selected fragments proved to be a rich source of structural insights. The simulations revealed that several fragments remained highly dynamic when bound to the Abu pocket. In the case of compound **3**, three different poses were characterised in details (Fig. 7, Fig. 8, Supplementary Fig. 3 and Supplementary Fig. 4). This may explain why the electron density maps of a number of fragments (e.g. **3**, **5** and **56**) were ambiguous and could not be fitted to a single binding pose. This suggests that it may be valuable to carry out MD simulations to assist model refinement process of X-ray diffracted crystals. The additional binding modes inferred from MD may also generate ideas for growing fragments out of the Abu pocket that may have been overlooked if design considerations had relied on a X-ray diffracted crystal structure showing a single pose. For instance the binding poses B and C of compound **3** suggest that Gly72 and Thr73 residues can form additional H-bonds interactions with Cyp binders.

### What did the fragments reveal about Cyclophilin inhibition strategies?

The binding modes of the various fragments to the Abu pocket of Cyp A were found to be remarkably similar to the interactions of compounds recently reported by the Pawlotsky and Guichou groups [38,46]. In these studies, several compounds were described as Cyp inhibitors and were co-crystallized in the active site of CypD. Gelin *et al.* and Ahmed-Belkacem *et al.* reported in particular that a para-aniline moiety was one of the very few substituents tolerated in the Abu pocket of CypD. The amino group forms hydrogen bonding interactions with Thr107 [38,46]. The X-ray structure of one inhibitor (PDB ID 4ZSC) was aligned with **3** from the present study (Supplementary Fig. 5). The fragment aligns well with the ligand and forms similar patterns of interactions (even though the isoforms differ). Considerable efforts have been invested by Ahmed-Belkacem *et al.* to replace the para-aniline moiety by an analogue that presents fewer toxicity liabilities for *in vivo* studies. The present results suggest that renewed chemistry efforts to merge the presently disclosed Abu pocket fragments with the scaffold from the lead series of Ahmed-Belkacem et al. may produce superior ligands for further drug development or *in vivo* investigations.

Finally, the three Cyp A Pro pocket fragments **16**, **60** and **61** (Fig. 5) were found to form interactions similar to those observed in crystal structures of Cyclosporine A in complex with Cyp A [10]. Specifically these three fragments are able to establish H-bond interactions with key structural and catalytic active site residues including Arg55, Gln63 and Asn102. This suggests that the binding modes of these fragments provide a good template for the elaboration of more potent lead-like molecules.

In conclusion, this work explored new ways of combining molecular dynamics simulations with biophysical measurement methods for fragment-based drug design. This work led to the discovery of several novel fragments that bind to Cyclophilin surfaces, and the resulting structural information will impact on future efforts to discover novel classes of potent and selective Cyclophilin ligands.

## Materials and Methods

### Protein expression purification and characterization

N-terminal hexa-histidine tagged human Cyclophilin (CypA, CypB and wt CypD) were provided by the Edinburgh Protein Production Facility (EPPF), University of Edinburgh. K133I-CypD plasmid was provided by Dr Jacqueline Dornan (Walkinshaw group, Edinburgh). Expression and purification for all isoforms was performed as described [42]. Dynamic light scattering (DLS) was used to verify the proteins’ quaternary structure and to examine any possible aggregation between Cyp monomers. DLS was performed in triplicates for each protein sample at 20 °C, on a Zetasizer APS instrument from Malvern using a 384 well plate with the total final volume of each well used to be 60μL. Prior DLS protein samples were buffer exchanged to PBS, concentrated to 1mg ml^−1^ and spun gently at 13,500 g for 20 minutes at 4 °C. Polydispersity index for all proteins was calculated as: 0.15 CypA, 0.16 CypB and 0.21 for CypD. Moreover, the hydrodynamic radius (*R*H) for each protein was also determined (4.3 ± 1.7 d.nm for CypA, 4.6 ± 1.8 d.nm for CypB and 4.3 ± 1.1 d.nm for CypD) and was in accordance with expected values for all Cyps, given existing X-ray crystallographic co-ordinates

### SPR

Compounds were initially screened at 1mM on a surface of 3200, 3000 and 3000 RU covalently stabilized His-CypA, -B and –D, respectively, in 10 mM PBS, 137 mM NaCl, 2.7 mM KCl, pH 7.4, 0.005 % v/v P20; 0.5 % v/v DMSO (2% v/v ethanol present to increase the solubility of some compounds), at 30 μl·min^−1^ with a 15 sec contact and dissociation time at 25°C. The surface was regenerated between each measurement and the compounds were washed off the surface of the sensor by running an excess of buffer solution at 40 μl.min^−1^ for 30 sec followed by a further 5 sec stabilization period. Selected compounds were further analysed with a 2-fold concentration series from 0.015 mM to 1 mM with similar solution conditions, flow rates and contact/dissociation time.

### Crystallization and data collection

Purified and his-tag cleaved CypA was concentrated to 30.4 mg ml^−1^ in PBS buffer. The vapor diffusion by hanging drop method at 6°C was used for the crystallization of CypA. The well volume was 1 mL, the precipitation solution consisted of 100 mM Tris-HCl, pH 8.0 and 21 – 24 % v/v PEG 8,000 and the drop consisted of 1.5 μL of CypA and 1.5 μL of well solution. Each ligand was soaked into an *apo* CypA crystal using 2 μl of soaking solution (30 % v/v PEG 8,000, 100 mM Tris-HCl,pH 8.0) and saturated concentrations of ligands (50 – 100 mM), followed by flash freezing with liquid nitrogen. Unliganded crystals were also prepared for obtaining *apo* CypA structures. X-ray intensity data were collected at the Diamond synchrotron-radiation facility in Oxfordshire, England. Intensity data for *apo* CypA and CypA-ligand complexes were collected from single crystals flash cooled in liquid nitrogen at 100 K. Data were processed with MOSFLM [47] and scaled with SCALA [48].

### Structure determination and refinement

All structures were originally processed with DIMPLE [49]. DIMPLE was followed by multiple cycles of restrained refinement using REFMAC 9 [50,51]. The side chains of the models were manually adjusted, while ligands and water molecules were added where appropriate using Coot 10 [52]. Further iterative cycles of restrained refinements and manual adjustments of ligands, side chains and water molecules, were carried out until the overall quality of the electron density maps could no longer be further improved.

### Preparation of proteins and ligands for free energy calculations

CypA, CypB and CypD structures were taken from the X-ray crystal structures with PDB IDs: 1CWA, 3ICH and 2BIT respectively. All water molecules were removed from the structures except the three tightly bound water molecules in the Abu pocket of cyclophilins [53]. All proteins were capped at the C-terminal and N-terminal with an N-methyl and acetyl groups, respectively. Protonation of the histidine residues was predicted by PROPKA, as implemented in Maestro Protein Preparation Wizard [54,55]. Specifically, the protonation states of CypA’s histidines were: HIE54, HIE70, HID92, HID126; in CypB: HIE54, HID92, HID126 and in CypD: HIE54, HIE70, HID92, HID126 and HID131. Ligands were prepared using Maestro as distributed by Schrödinger [54]. A number of fragments were expected to be almost exclusively charged in assay conditions (pH 7.4-8.0). Since correcting for net-charge changes is non-trivial in free energy calculations, these compounds were not included in the dataset for free energy calculations. [56,57] On the basis of pKa calculations a number of the remaining fragments could potentially adopt different protonation states in solution and/or bound to the protein. For similar reasons only the neutral form was considered in subsequent docking and free energy calculations.

### Docking calculations

All ligands were docked into the Abu and Pro pockets of CypA, CypB and CypD, in order to generate starting conformations for the free energy calculations, using the Vina software via the Autodock Vina plugin of Pymol [58–60]. For the docking calculations in the Abu pocket, the box size was set to *x* = *y* = *z* = 13.12 Å and the box was centered on the Abu. The whole protein was used as a receptor and, unless otherwise mentioned, only the top-ranked pose was retained for each ligand. Moreover, three water molecules that are tightly bound in Abu pocket [53] were kept and treated as part of the protein for docking purposes. All ligands were docked in the Pro pocket of CypA, CypB and CypD, in a similar way. The box size was set to *x* = *y* = *z* = 7.50 Å and was centered on the Pro pocket.

### Free energy calculations set up

The strategy used here consisted in carrying out series of relative free energy calculations to connect all compounds in the dataset to a reference compound, and then to compute a standard (absolute) binding free energy for the reference compound. This two-step procedure yields standard binding free energies for all compounds in the dataset.

For the calculation of relative binding free energies, ligands should be perturbed from one ligand to another in complex with the protein and alone in a water box. For this reason, a perturbation map was generated (Supplementary Fig. 6), by manually connecting all the ligands via multiple transformations. Ligand **3** was set as the center of the map, due to its high chemical similarity to the rest of the compounds. All ligands were connected by keeping the number of perturbation steps to **3** to less than four. Mappings featured compounds with high structural similarity, in order to reduce the number of perturbed atoms in each relative free energy calculation. Input files for all subsequent free energy simulations were set up using FESetup1.1 software [61].

Protein parametrization was done using the Amber ff14SB force field [62], while ligands were parametrized using the GAFF force field as implemented in Amber14 [63,64] and AM1-BCC charges [65,66]. All systems were solubilized in a rectangular box of TIP3P water molecules, with a box length of 10 Å away from the edge of the solute, and Na^+^ or Cl^−^ ions were added to neutralize the net charge of the system. The systems were energy minimized for 200 steps to alleviate any steric clashes, followed by a heating step to 300 K for 200 ps, with harmonic potential restraints on all non-solvent atoms using a 10 kcal mol^−1^ Å^2^ force constant. Systems were then equilibrated by running short (200 ps) MD runs using NVT ensemble with the same restraints as before. Finally, to stabilize the density a final run of 5 ns using an NPT ensemble at 1 atm was performed. The last trajectory snapshot was used as the starting point for the free energy calculations.

### Alchemical free energy calculations

Relative free energy changes for transforming a ligand L1 into a ligand L2 (*ΔΔG*_(L1→L2)_) were calculated as the difference in the free energy change of transforming L1 to L2 in a water box (*ΔG_w_*(L1→L2)) and in complex with the protein (*ΔG_p_*_(L1→L2)_) as shown by equation1:

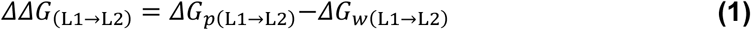

The MBAR implementation in the software pymbar was used for the calculation of both *ΔG_w_*_(L1→L2)_ and *ΔG_p_*_(L1→L2)_ free energy changes [67]. Relative binding free energy calculations were only performed once, and statistical errors were propagated from each step, with the final reported error calculated using equation 2

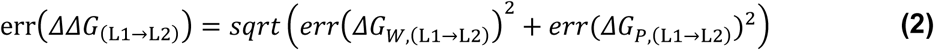

where the terms on the right hand side of eq. 2 are the statistical errors associated with *ΔG_w_*_(L1→L2)_ and *ΔG_p_*_(L1→L2)_ respectively.

To assess the sensitivity of the calculated relative free energies to the starting conditions of the simulations, the calculations were repeated three times for a subset of 10 fragments (**1**, **2**, **4**, **5**, **7**, **42**, **49**, **82**, **96** and **97** as indicated in Supplementary Fig. 6). Each repeat was setup from the first, second and third top-ranked poses produced by vina, and initial velocities were drawn from the Maxwell-Boltzmann distribution. For this dataset the mean unsigned difference between the average relative binding free energy estimated from the 3 repeats and from a single repeat is 0.35 kcal.mol-1, and the mean standard deviation is 0.5 kcal.mol-1 (See Supplementary Dataset S1). Thus the precision of the calculated binding free energies obtained from a single repeat may be considered to be ca. 0.5 kcal.mol-1. This figure indicates uncertainties typically higher than what was obtained by use of equation 2, but was deemed sufficient to discriminate Abu/Pro pocket binding preferences, and strong/weak binders within each pocket (Figure 2).

The absolute binding free energy of ligand L3 was calculated with equation 3:

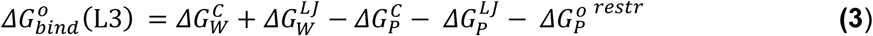

where 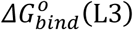 is the absolute binding free energy of ligand **3** (L3). 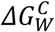 and 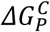 are the free energies of turning off the partial charges of L3 in water and in complex respectively, while 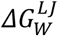 and 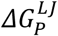 are the free energies of turning off the Lenard Jones parameters of the uncharged ligand. 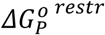 is the free energy cost of releasing the restraints to reach standard state conditions, as explained in ref [68]. This quantity was computed with a custom script as described in ref [44]. Each absolute binding free energy calculation was repeated in quadruplicate and the reported value is the average of the quadruplicate. The final error reported for the absolute binding free energy was calculated as the standard error of the mean of the four repeats.

Free energy simulations were performed using the SOMD (Sire – OpenMM) framework [69,70] on GPUs (GeForce GTX465 and Tesla/M2090/K20 graphic cards). Twelve equidistant λ windows (0.0000, 0.0909, 0.1818, 0.2727, 0.3636, 0.4545, 0.5455, 0.6364, 0.7372, 0.8182, 0.9091, 1.0000) were used and the systems were energy minimized for 1000 steps and then re-equilibrated at the appropriate λ value for 20 ps. The total length of each simulation (at each λ) was 5 ns, and the perturbed energies were saved every 200 fs. The perturbed energies where post-processed using the MBAR estimator [71]. A softcore potential was used in all simulations [72]. The hydrogen mass repartitioning (HMR) technique [73] was used to increase the timestep of the simulation and decrease the time needed to run each simulation. A 4 fs timestep and a repartitioning factor of 4 were used and all bonds were constrained. Simulations were performed in the NPT ensemble, where temperature control was achieved by using the Andersen thermostat, [74] and pressure control was achieved by a Monte Carlo Barostat. Periodic boundary conditions were applied with a 10 Å cut-off for the non-bonded interactions.

In relative binding free energy simulations interatomic flat-bottom distance restraints of 7.5 ± 3.0 Å in the Abu and Pro pockets were applied with a force constant of 5 kcal mol^−1^. Flat-bottom distance restraints allow the fragments to move freely in the pocket as long as the specified distance is within 4.5 and 10.5 Å. If the specified distance is below 4.5 or above 10.5 Å then a force constant of 5 kcal mol^−1^ is applied to bring this distance back to the allowed region. One atom of each ligand was selected and restrained to one protein atom in the Abu pocket (Gln111/backbone O atom) in the case of ligands present in the Abu pocket. For the simulations of ligands in the Pro pocket, the protein atom was Met61/CA and the ligand atom was the same as in the Abu pocket. Although restraints are not usually needed in relative binding free energy simulations, these were considered necessary in the present simulations. This is because fragments weakly bound to a shallow pocket can drift away from their starting binding position into bulk after a few ns of simulation, leading to the evaluation of free energy changes that no longer correspond to interactions with the desired pocket. The selected atomic distance flat bottom restraints (7.5 ± 3.0 Å) and the force constant (5 kcal mol^−1^) were sufficient to restrain the fragments in the binding pocket and prevent them from drifting away, while at the same time, avoiding unduly penalizing motions in the Abu or Pro pockets.

For reference compound **3,** absolute binding free energy calculations were performed to both the Abu and Pro pockets of CypA. Ligand **3** was restrained in the pocket of the protein using interatomic flat-bottom distance restraints with a force constant of 5 kcal mol^−1^. One atom of ligand **3** was distance restrained to two atoms of the protein in the Abu pocket (GLN111 and ASN102 CA backbone atoms) or to two atoms of the protein in the Pro pocket (HID126 and ILE57 CB atoms). In the Abu pocket the distance restraint tolerance was set to 5.5 ± 1.5 Å and in the Pro pocket to 7.5 ± 1.0 Å. The free energy cost for removing these restraints was evaluated by post-processing the simulation trajectories to yield a standard free energy of binding [44].

Supplementary dataset S1 contains all input files for the FESetup and SOMD codes, and a summary of the calculated free energies obtained by post-processing of the generated trajectories. The dataset can be downloaded from the github repository located at https://github.com/michellab/cypafragmentsfreeenergy.

## Accession numbers

Coordinates and structure factors for all ligands binding CypA were deposited in the protein data bank with accession numbers: CypA_lig3 (**5NOQ**), CypA_lig5 (**5NOR**), CypA_lig16 (**5NOS**), CypA_lig56 (**5NOT**), CypA_lig60 (**5NOU**), CypA_lig61 (**5NOV**), CypA_lig89 (**5NOW**), CypA_lig97 (**5NOX**), CypA_lig98 (**5NOY**), CypA_lig99 (**5NOZ**).

## Acknowledgements

Gratitude is expressed to BBSRC and the EaStBio DTP for funding this project. Julien Michel is supported by a University Research Fellowship from the Royal Society. The research leading to these results has received funding from the European Research Council under the European Union’s Seventh Framework Programme (FP7/2007-2013)/ERC grant agreement No. 336289.

## Abbreviations

Cyps: Cyclophilins
CypA: cyclophilin A
PPIases: peptidyl-prolyl isomerases
CsA: cyclosporin A
HIV-1: human immunodeficiency virus 1
HCV: hepatitis C virus
SBDD: structure-based drug design
FBDD: fragment based drug discovery
MD: molecular dynamics
SPR: surface plasmon resonance
HBD: hydrogen bond donor
HBA: hydrogen bond acceptor
DMSO: dimethyl sulfoxide
PBS: phosphate buffered saline

